# A network pharmacology approach to assess the comparative pharmacodynamics of pharmaceutical excipient trehalose in human, mouse and rat

**DOI:** 10.1101/2023.01.23.525154

**Authors:** Jack Friend, Arun HS Kumar

## Abstract

**Background:** Trehalose is used as a pharmaceutical excipient due to its several desirable pharmacokinetic and historically evident safety features. However, information on the pharmacodynamic properties of trehalose is lacking. Hence this study evaluated the comparative pharmacodynamic properties of trehalose using a network pharmacology approach.

**Materials and methods:** The specific targets of trehalose in human, mouse and rat were identified from the SwissTargetPrediction database, categorised and compared. The expression profile and subcellular localisation of the targets of trehalose in human was identified and correlated with the affinity of trehalose to these targets to assess its impact on the pharmacodynamic properties of trehalose. The affinity of trehalose to its metabolising enzyme in human, mouse, and rat was assessed by molecular docking and compared.

**Results:** A significant difference in the target categories and target types of trehalose was observed in human, mouse, and rat. The affinity of trehalose to human (66.03 ± 5.1 μM), rat (102.53 ± 11.3 μM) and mouse (42.07 ± 5.3 μM) trehalase was significantly different. Family A G protein coupled receptors were identified as the major target category of trehalose and cyclin dependent kinase 1 was observed as the high affinity target of trehalose in human and mouse. The correlation of target expression and affinity indicated minimal pharmacodynamic influence under physiological conditions.

**Conclusion:** This study reports the selective targets of trehalose relevant to drug discovery and development protocols and highlights the limitations of rodent models in translating pharmacodynamic analysis of trehalose for development of human therapeutics.

## Introduction

Pharmaceutical excipients are ideally anticipated to be pharmacodynamically inert, and they are used to formulate, stabilise, deliver, and preserve the active pharmaceutical ingredient (API). However, like the API, excipients may also have their own bioactivity.^[1–4]^ Many excipients used in pharmaceutical formulations are Generally Recognized as Safe (GRAS) compounds in the food industry, and the trust in these compounds may be conferred by their long history of use in consumer products and various pharmaceutical formulations.^[5–9]^ In both Europe and the United States, formulations with commercially established excipients may be granted approval without the need for the full, excipient specific toxicology report that is required for new chemical entities. Similarly, excipients with prior use in Japan are not evaluated at the same level as novel excipients without prior use.^[10]^

Trehalose is a disaccharide that is used in oral and intravenous pharmaceutical formulations. It is a nonreducing disaccharide formed from two α-glucose joined by a 1-1 glycosidic bond. The alpha-1,1-isomer is quite common across fungi, plants, and insects where it is suggested that trehalose is used as a natural energy storehouse or cryptobiotic.^[11, 12]^ Trehalose synthesis is absent in vertebrates, and there are no trehalose synthesis genes in vertebrates.^[13]^ However, the trehalose catabolizing enzyme trehalase (EC 3.2.1.28) is reported in vertebrates, including mammals, where it is anchored to the brush border membrane of the intestine and kidney epithelium.^[14]^ The nonreducing alpha-1,1-glycosidic bond makes trehalose an exceptionally stable sugar that can withstand acid hydrolysis and cleavage by alpha--glycosidase. Crystalline dihydrate trehalose does not readily absorb water, and it is stable up to 97°C. Compared to sucrose (another disaccharide with similar molecular weight) trehalose has a lower solubility, a lower diffusion coefficient, a higher hydration number, and a higher viscosity. Trehalose can stabilise protein or lipid preparations through direct interactions by hydrogen bonding to the hydrophilic protein surface, or indirectly by binding water.^[6–9, 15]^

The desirable physical properties make trehalose an ideal excipient in drug formulations, and hence it has been used in approved medicinal products, such as the vaccines for ovine rinderpest, and mammalian cell culture desiccation.^[16, 17]^ Recently, trehalose is also used as an excipient for formulating monoclonal antibodies (Genetech’s Herceptin, Avastin, and Lucentis), and recombinant protein (Baxter’s Advate).^[15]^ Given that trehalose’s GRAS status is based on its historical use as a food additive (GRAS Notice No. 912), it is important to study the bioactivity of trehalose, especially when it bypasses the intestine through intravenous and intramuscular injections. When trehalose is used in oral formulations, it is hydrolysed by trehalase and the potential impact of this metabolism on gastrointestinal physiology needs critical assessment.

Rodent models, especially mice, are the most common animal used in biomedical research studies. While the use of rodent models has been indispensable, we have found that these models do not always translate well to humans.^[18]^ This may be due unanticipated to physiological differences.^[19, 20]^ Hence in this study using *in silico* methods, we reverse screened the protein targets of trehalose in *Homo sapiens*, *Mus musculus*, and *Rattus norvegicus* (hereafter human, mouse, and rat) for a comparative and evidence based assessment of its network pharmacology and pharmacodynamic properties.

## Materials and Methods

The SMILES structure of trehalose acquired from the PubChem database was inputted into SwissTargetPrediction database to identify was human, mouse, and rat specific targets.^[21–25]^ The target list of trehalose was processed based on their probability scores to identify highest affinity target/s and only the targets with a probability score greater than 0 were used for further analysis (**Supplement S1**). Enzyme trehalase (EC 3.2.1.28) was identified as a binding target by SwissTargetPrediction, but with 0 probability of binding but only being influenced by its enzymatic activity, hence it was excluded as a binding target. The SwissADME database was used to assess the physiochemical and pharmacokinetic parameters of Trehalose (**Supplement S1**).^[21]^ The targets of trehalose were subclassified into functional categories and the relative proportion of each of the functional categories among the total number of targets was estimated and compared between human, mouse, and rat.

We used the R package HPAanalyze to access tissue and subcellular expression (protein and mRNA) of trehalose targets in humans from the Human Protein Atlas (**Supplement S1**).^[22, 26–28]^ The data for protein, mRNA and subcellular expression were graphed using ggplot2.^[29]^ The protein expression was profiled under following four categories: high, medium, low and no expression. The mRNA expression was categorised based on protein transcripts per kilobase million (pTPM) ranging from 0 to 1250. While the subcellular locations of the trehalose targets were dichotomized into being absent or present. The expression data of trehalase protein in human tissues was also assessed and categorised into high, medium, or low expression (**Supplement S1**). The affinity of trehalose to human, mouse, and rat specific trehalase was assessed by molecular docking using AutoDock vina 1.2.0 as reported previously.^[25, 30, 31]^

## Results

Trehalose is used in oral, intravenous, and intramuscular formulations due to its wider and compatible pharmacokinetic properties (**table 1**). The low lipophilicity (iLOGP = 0.57, consensus Log P = −3.55) and high water solubility (ESOL Log S = 0.94) of trehalose supports its use in a diverse range of formulation types. The pharmacokinetic parameters suggest that trehalose cannot easily cross the plasma membrane unless facilitated by a specific or selective transporter. Predictably, trehalose has a low bioavailability (0.17), low gastrointestinal absorption, and it is impermeable to the Blood-Brain Barrier and skin barrier (log Kp = −11.36 cm/s). Trehalose was also an observed substrate for p-glycoprotein which further decreases its gastrointestinal absorption. Trehalose did not appear to inhibit cytochrome liver enzymes (CYP1A2, CYP2C19, CYP2C9, CYP2D6, and CYP3A4) assessed in this study, suggesting a favourable feature as an excipient that will not affect the pharmacokinetics of the API in the formulation.

**Table 1.**
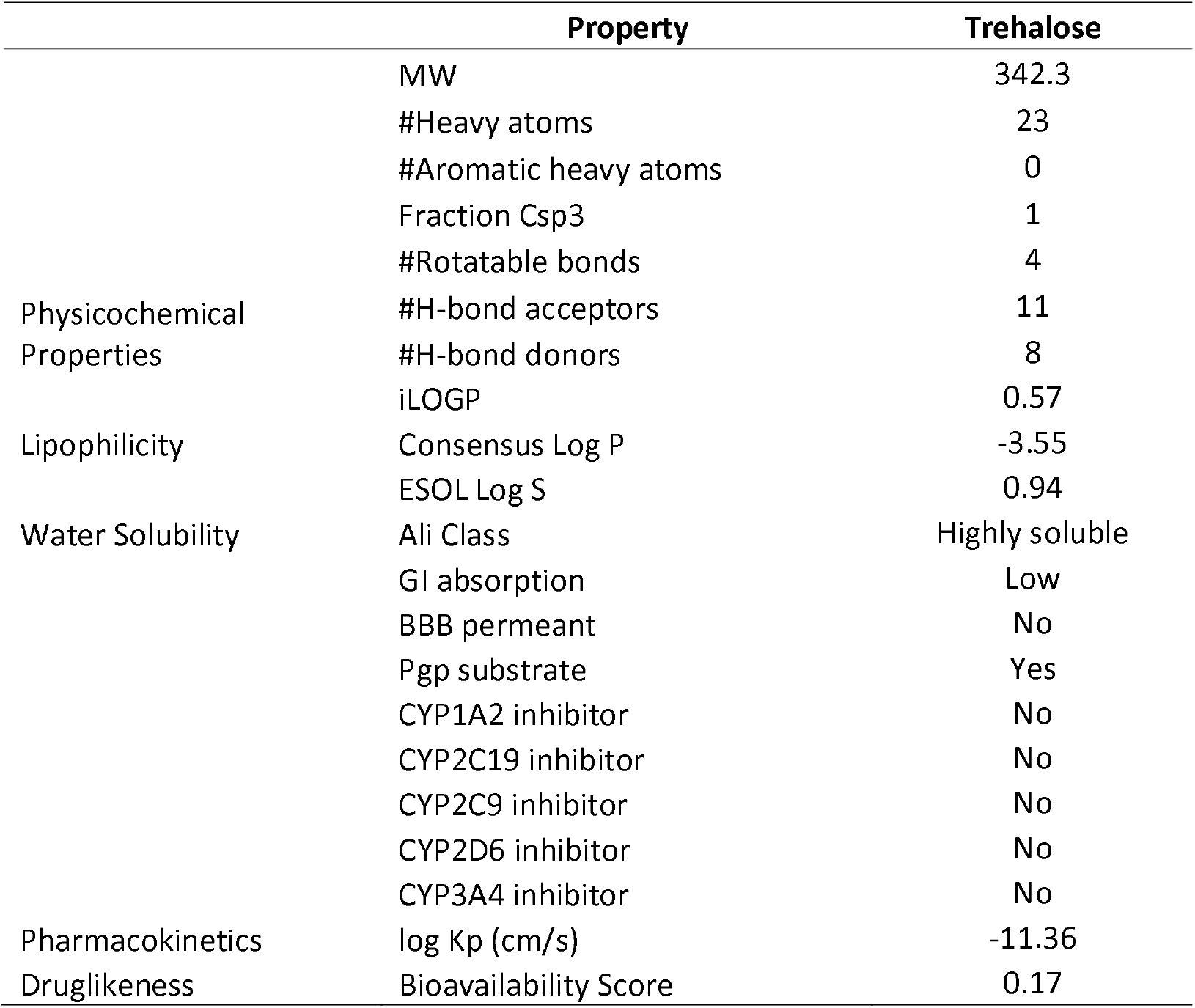
Physicochemical and Pharmacokinetic properties of trehalose.

Trehalose is metabolised by trehalase in mammals, therefore tissue specific expression of this enzyme in human was assessed. Trehalase is moderately expressed in kidney, duodenum, and small intestine, while its expression in ovary, adrenal gland, colon and cerebral cortex was low (**table 2**). This suggests that trehalose will be preferentially metabolised in the kidneys, duodenum, and small intestine, so it may not be a preferred excipient if the intention is to deliver the API to these target organs. We also observed considerable variability in the affinity of trehalose to human, mouse, and rat trehalase. Mouse trehalase had the highest affinity for trehalase, followed by human trehalase, and least of all rat trehalase (**table 2**). This significant species specific variability in enzyme-substrate affinity becomes important in drug discovery studies. Hence considerable caution is necessary when extrapolating findings from preclinical rodent studies to design phase 1 clinical trials involving human subjects.

**Table 2.**
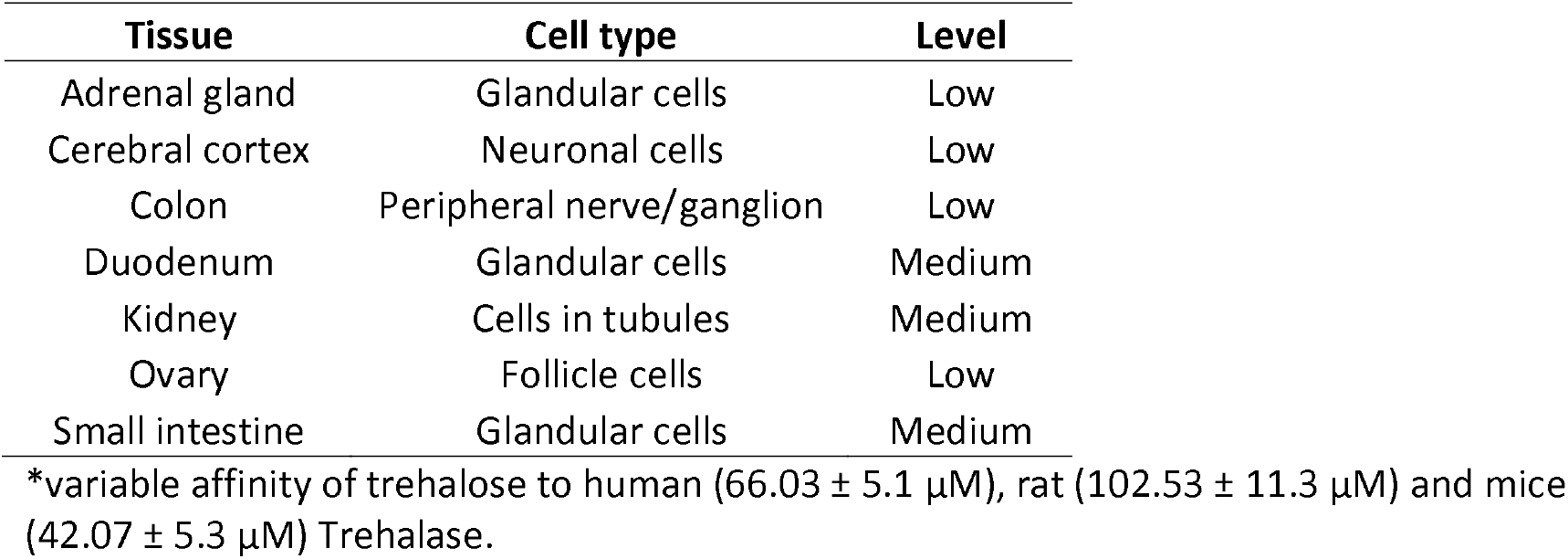
Protein level expression of trehalase enzyme in human and the affinity of trehalose to human, rat and mice specific trehalase.

| Tissue | Cell type | Level |
| --- | --- | --- |
| Adrenal gland | Glandular cells | Low |
| Cerebral cortex | Neuronal cells | Low |
| Colon | Peripheral nerve/ganglion | Low |
| Duodenum | Glandular cells | Medium |
| Kidney | Cells in tubules | Medium |
| Ovary | Follicle cells | Low |
| Small intestine | Glandular cells | Medium |

\*variable affinity of trehalose to human ( $66.03 \pm 5.1 \mu\text{M}$ ), rat ( $102.53 \pm 11.3 \mu\text{M}$ ) and mice ( $42.07 \pm 5.3 \mu\text{M}$ ) Trehalase.

The species specific variability of trehalose was further evident from differences in its binding targets in human, mouse, and rat. Family A G Protein Coupled Receptors (GPCRs) made up the largest portion of binding target classes of trehalose. This was conserved across human (43%), mouse (43%), and rat (40%) (**figure 1**). However, the nature of the other target classes of trehalose differed considerably between the three species. In human, the next three most prominent target classes were enzymes (13%), other cytosolic proteins (13%), and secreted proteins (10%). Other cytosolic proteins were also identified as a prominent target class in mouse (14%), and enzymes were identified as a prominent target class in rat (20%). Secreted proteins were not identified as targets in either mouse or rat. In mouse, the remaining target classes were kinases (14%), nuclear receptors (14%), and transcription factors (14%). This was not conserved between human and mouse. Kinases, nuclear receptors, and transcription factors each made up 3% of binding targets in human. In rat, the remaining target classes were kinases (10%), ligand-gated ion channels (10%), nuclear receptors (10%), or unclassified proteins (10%). Unclassified proteins were not identified as binding targets in human. This data suggests that protein binding targets largely differ between humans and rodent models. Therefore, preclinical safety and efficacy assessments of trehalose using rodent models may not be relevant for extrapolating the findings to design human clinical trials.

**Figure 1:**
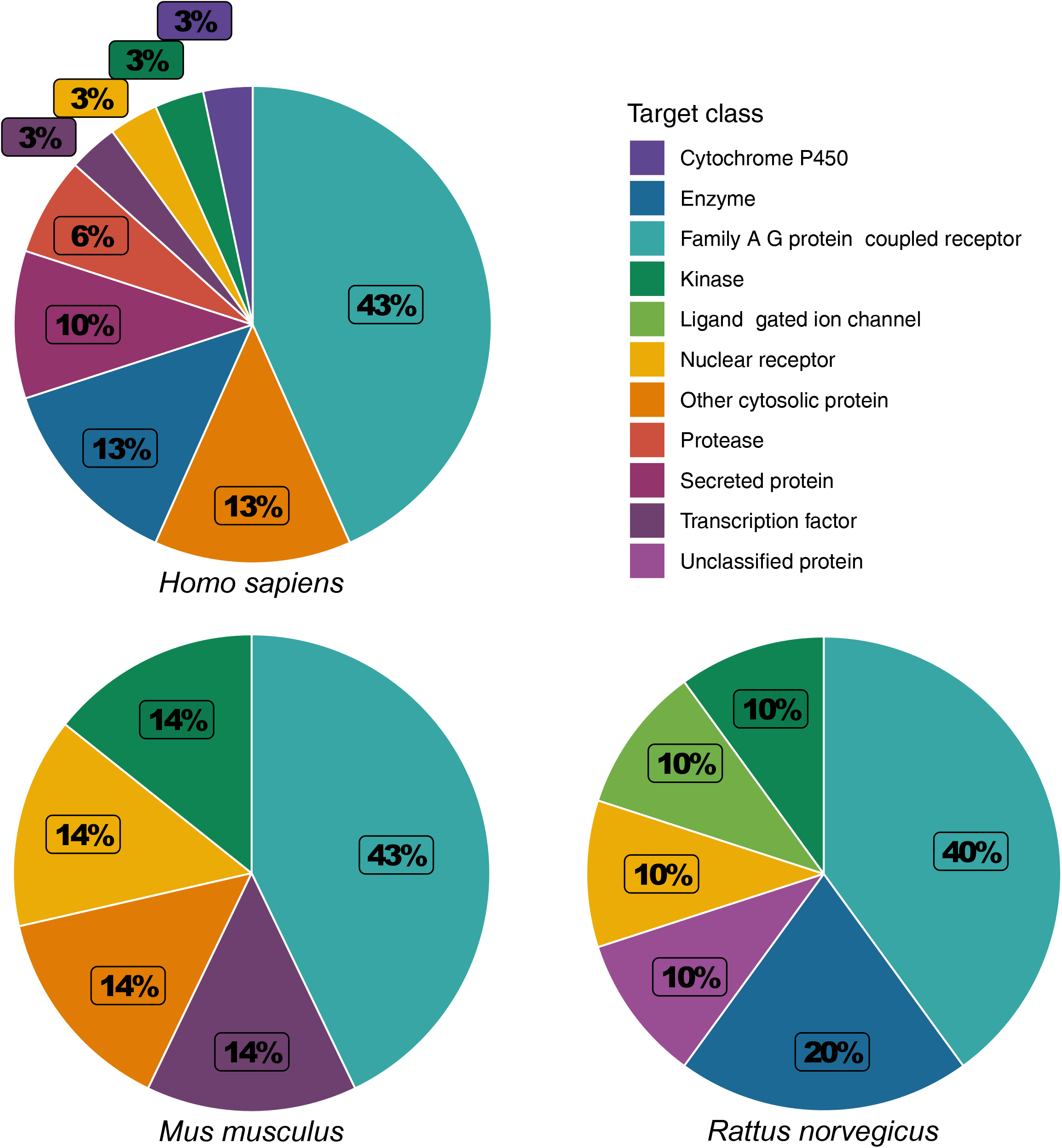
The target categories (target class) of predicted trehalose targets in human, mouse, and rat are represented as percentages.

Variations in the binding targets of trehalose were observed between human, mouse and rat (**table 3**), once again highlighting potential irrelevance of rodent models in assessing trehalose containing formulations for therapeutic applications in human. Cyclin-dependent kinase 1 (CDK1) is serine/threonine protein kinase involved in cell cycle regulation and transcription at the G1 checkpoint,^[32]^ and it had the strongest binding affinity for trehalose (probability score = 0.24). CDK1 and heat shock protein HSP 90-alpha (HSP90AA1) were predicted binding targets of trehalose in both human and mouse, but not in rat. Dopamine D2 receptor (DRD2) was a predicted binding targets of trehalose in human, mouse, and rat with similar low affinity. Dopamine D1 receptor (DRD1) was observed in only human and mouse. Dopamine D3 receptor (DRD3) was observed in only human and rat. While all serotonin (5-HT) receptors were predicted targets of trehalose in human, mouse, and rat, different receptors and subunits are targeted. Serotonin 1 receptor was observed in both human (HTR1B) and rat (HTR1A and HTR1B), but not mouse. Serotonin 2 receptor was observed in human (HTR2A, HTR2B, and HTR2C), mouse (HTR2C), and rat (HTR2A). Serotonin 6 receptor (HTR6) was only observed in human. RAR-related orphan receptor gamma (RORC) and STAT3 were predicted binding targets of trehalose in human and mouse, but not in rat. Alpha adrenergic receptors were predicted binding targets of trehalose in human and rat, but not in mouse. The alpha 1a adrenergic receptor (ADRA1A) was observed in both human and rat. Alpha 1b adrenergic receptor (ADRA1B) was observed in only rat, and alpha 1d adrenergic receptor (ADRA1D) was observed in only human. The alpha 2b and 2c adrenergic receptors (ADRA2B and ADRA2C) were observed in both human and rat. Glutamate carboxypeptidase II (FOLH1) was a predicted binding target of trehalose in human and rat, but not mouse.

**Table 3.** The binding targets of trehalose in *Homo sapiens*, *Mus musculus*, and *Rattus norvegicus*.

|  | Target protein | Gene | Target protein class | Probability |
| --- | --- | --- | --- | --- |
| *Homo sapiens* | Cyclin-dependent kinase 1 | CDK1 | Kinase | 0.24 |
|  | Heat shock protein HSP 90-alpha | HSP90AA1 | Other cytosolic protein | 0.11 |
|  | Vascular endothelial growth factor A | VEGFA | Secreted protein | 0.11 |
|  | Gamma-secretase | PSEN2 | Protease | 0.11 |
|  | Gamma-secretase | PSENEN | Protease | 0.11 |
|  | Gamma-secretase | NCSTN | Protease | 0.11 |
|  | Gamma-secretase | APH1A | Protease | 0.11 |
|  | Gamma-secretase | PSEN1 | Protease | 0.11 |
|  | Gamma-secretase | APH1B | Protease | 0.11 |
|  | Acidic fibroblast growth factor | FGF1 | Secreted protein | 0.11 |
|  | Heparanase | HPSE | Enzyme | 0.11 |
|  | Basic fibroblast growth factor | FGF2 | Secreted protein | 0.10 |
|  | Galectin-4 | LGALS4 | Other cytosolic protein | 0.10 |
|  | Galectin-3 | LGALS3 | Other cytosolic protein | 0.10 |
|  | Galectin-8 | LGALS8 | Other cytosolic protein | 0.10 |
|  | Glutamate carboxypeptidase II | FOLH1 | Protease | 0.10 |
|  | Serotonin 2b (5-HT2b) receptor | HTR2B | GPCR (Family A) | 0.10 |
|  | Alpha-2a adrenergic receptor | ADRA2A | GPCR (Family A) | 0.10 |
|  | Adrenergic receptor alpha-2 | ADRA2C | GPCR (Family A) | 0.10 |
|  | Alpha-2b adrenergic receptor | ADRA2B | GPCR (Family A) | 0.10 |
|  | Dopamine D1 receptor | DRD1 | GPCR (Family A) | 0.10 |
|  | Dopamine D2 receptor | DRD2 | GPCR (Family A) | 0.10 |
|  | Alpha-1d adrenergic receptor | ADRA1D | GPCR (Family A) | 0.10 |
|  | Serotonin 2a (5-HT2a) receptor | HTR2A | GPCR (Family A) | 0.10 |
|  | Serotonin 2c (5-HT2c) receptor | HTR2C | GPCR (Family A) | 0.10 |
|  | Dopamine D3 receptor | DRD3 | GPCR (Family A) | 0.10 |
|  | Cytochrome P450 2D6 | CYP2D6 | Cytochrome P450 | 0.10 |
|  | Serotonin 6 (5-HT6) receptor | HTR6 | GPCR (Family A) | 0.10 |
|  | Alpha-1a adrenergic receptor * | ADRA1A | GPCR (Family A) | 0.10 |
|  | Serotonin 1b (5-HT1b) receptor * | HTR1B | GPCR (Family A) | 0.10 |
|  | Nuclear receptor ROR-gamma | RORC | Nuclear receptor | 0.10 |
|  | Signal transducer and activator of transcription 3 | STAT3 | Transcription factor | 0.10 |
|  | Sialidase 3 | NEU3 | Enzyme | 0.10 |
|  | Sialidase 2 | NEU2 | Enzyme | 0.10 |
|  | Sialidase 4 | NEU4 | Enzyme | 0.10 |
| *Mus musculus* | Cyclin-dependent kinase 1 * | Cdk1 | Kinase | 0.24 |
|  | Heat shock protein HSP 90-alpha * | Hsp90aa1 | Other cytosolic protein | 0.11 |
|  | Dopamine D1 receptor * | Drd1 | GPCR (Family A) | 0.10 |
|  | Dopamine D2 receptor * | Drd2 | GPCR (Family A) | 0.10 |
|  | Serotonin 2c (5-HT2c) receptor * | Htr2c | GPCR (Family A) | 0.10 |
|  | Nuclear receptor ROR-gamma * | Rorc | Nuclear receptor | 0.10 |
|  | Signal transducer and activator of transcription 3 * | Stat3 | Transcription factor | 0.10 |
| *Rattus norvegicus* | Glutamate carboxypeptidase II | Folh1 | Enzyme | 0.10 |
|  | Serotonin 1a (5-HT1a) receptor | Htr1a | GPCR (Family A) | 0.10 |
|  | Alpha-1b adrenergic receptor | Adra1b | GPCR (Family A) | 0.10 |
|  | Alpha-1a adrenergic receptor | Adra1a | Ligand-gated ion channel | 0.10 |
|  | Serotonin 1b (5-HT1b) receptor | Htr1b | GPCR (Family A) | 0.10 |
|  | Alpha-2c adrenergic receptor * | Adra2c | Nuclear receptor | 0.10 |
|  | Alpha-2b adrenergic receptor * | Adra2b | Unclassified protein | 0.10 |
|  | Dopamine D2 receptor * | Drd2 | GPCR (Family A) | 0.10 |
|  | Serotonin 2a (5-HT2a) receptor * | Htr2a | Kinase | 0.10 |
|  | Dopamine D3 receptor * | Drd3 | Enzyme | 0.10 |

Performed with SwissTargetPrediction. \* = predicted by homology. GPCR = G protein coupled receptor.

We assessed the protein expression profile of the targets of trehalose in human. The protein expression of trehalose targets were categorised into four categories – high, medium, low, and no expression (**figure 2**). Presenilin is a membrane bound protease that makes up the catalytic unit of the gamma secretase complex, and together they may cleave amyloid precursor protein into beta-amyloid.^[33]^ Generally, presenilin/gamma secretase associated proteins had a high, widely distributed expression. The alpha-1 homolog A (APH1A) subunit had the highest expression across all body tissues, and it was at least moderately expressed in all tissues except for skeletal muscle, ovary, and the cerebral cortex where it only had low expression. Presenilin-1 (PSEN1) was widely expressed, but absent from adipose and oral mucosa. In contrast, presenilin enhancer protein 2 (PSENEN) was absent from the parathyroid gland, lung, and testis. Nicastrin (NCSTN) was absent from adipose tissue, heart muscle, smooth muscle, oral mucosa, skeletal muscle, bone marrow, soft tissue, lymph node, and spleen. At least three of the four observed presenilin/gamma secretase proteins were present in every tissue, except for adipose tissue, and oral mucosa. There was also ubiquitously low expression of these presenilin/gamma secretase associated proteins in the ovary.

**Figure 2:**
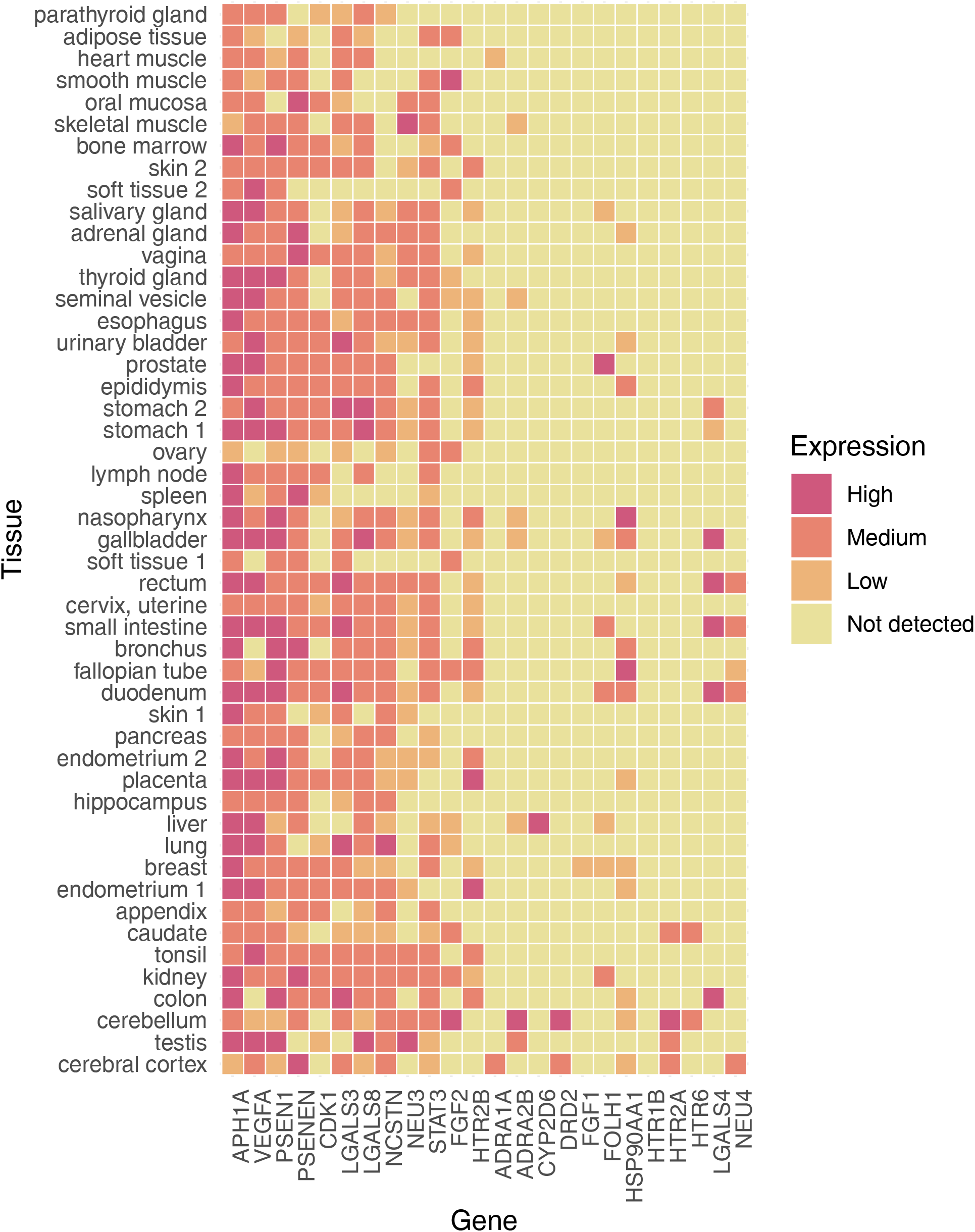
The protein expression profile of predicted trehalose targets in human. The expression profile is categorised into the following high (red), medium (orange), low (light orange) and no expression (yellow).

We observed three families of GPCR neurotransmitter receptors: the serotonin (5-HT) receptor family, the alpha adrenergic receptors, and the dopamine receptor family (**figure 2**). Serotonin 6 (HTR6) receptor had medium expression in the caudate nucleus and cerebellum. Serotonin 2a receptor (HTR2A) was also expressed in the cerebellum and caudate, but also the testis and cerebral cortex. Serotonin 2b receptor (HTR2B) was widely expressed throughout skin, glands, and gastrointestinal tract, and female reproductive tract. Alpha-1a adrenergic receptor (ADRA1A) was only expressed in heart muscle and cerebral cortex. Alpha-2b adrenergic receptor (ADRA2A) was highly expressed in the cerebellum, and moderately/lowly expressed in skeletal muscle, seminal gland, nasopharynx, gallbladder, liver and testis. Dopamine D2 receptor (DRD2) was highly expressed in the cerebellum and moderately expressed in cerebral cortex.

Galectins are a family of alpha-galactoside carbohydrate binding proteins.^[34]^ Galectin-3 (LGALS3), galectin-4 (LGALS4), and galectin-8 (LGALS8) were observed (**figure 2**). Galectin-4 was highly expressed in a few select tissues (gallbladder, rectum, small intestine, duodenum, and colon). Galectin-3 and galectin-8 had similar expression patterns where both were absent from soft tissue and spleen. Between the two, they had either high or medium expression in the urinary bladder, stomach, gallbladder, rectum, small intestine, duodenum, lung, and colon – a pattern which overlaps with galectin-4. Notably, galactin-8 was highly expressed in the testis, but galectin-4 and galactin-3 were absent.

We observed salidase-3 (NEU3) and salidase-4 (NEU4), hydrolases that cleave the glycosidic bond in neuraminic acids (**figure 2**). Viral neuraminidase, a drug target for the prevention of the spread of influenza infection.^[35]^ Salidase-3 is highly expressed in skeletal muscle and testis. Salidase-4 is moderately expressed in the rectum, small intestine, duodenum, and cerebral cortex, and has low expression in the fallopian tube. Salidase-3 is absent from the cerebral cortex and hippocampus, but has moderate expression in the cerebellum. On the other hand, salidase-4 has moderate expression in the cerebral cortex, and it is absent from the cerebellum and hippocampus.

CDK1, which was the highest binding affinity target of trehalose in human and mouse (**table 3**), has moderate expression across body tissues. Since CDK1 is involved in driving the cell cycle and cell proliferation,^[32]^ administration of trehalose containing formulations may affect these tissues. Vascular endothelial growth factor A (VEGFA) is glycoprotein associated with neurons that promotes angiogenesis.^[36]^ VEGFA was widely expressed except for ovary, bronchus, and colon. It was highly expressed across the reproductive organs, GI tract, liver, and lung. It was only moderately expressed in the cerebral cortex, cerebellum, and hippocampus. STAT3 is a transcription factor that is activated by the Janus kinases in the JAK/STAT pathway.^[37]^ STAT3 had either medium or low expression in the nervous system, muscle, glands, and gastrointestinal tract. Glutamate carboxypeptidase II (FOLH1) was expressed in the kidney, breast, liver, duodenum, small intestine, gallbladder, prostate, and salivary gland. HSP90AA1 was highly expressed in the fallopian tube and nasopharynx. Overall the cerebral cortex, testis, and cerebellum tissues had the highest composite expression of binding targets of trehalose in human, followed by target concentration in the colon and kidney (**figure 2**). We were not able to obtain protein expression data for eleven proteins: several alpha adrenergic receptors (ADRA1D, ADRA2A, ADRA2C), two dopamine D receptors (DRD1, DRD3), presenilin/gamma secretase (APH1B and PSEN2), a serotonin 2c receptor (HTR2C), sialidase 2 (NEU2), Nuclear receptor ROR-gamma (RORC), and Heparanase (HPSE). RNA transcript expression of all selected genes were graphed. Unsurprisingly, protein expression and RNA transcript expression data largely did not overlap for the available data (**figures 2 and 3**).

**Figure 3:**
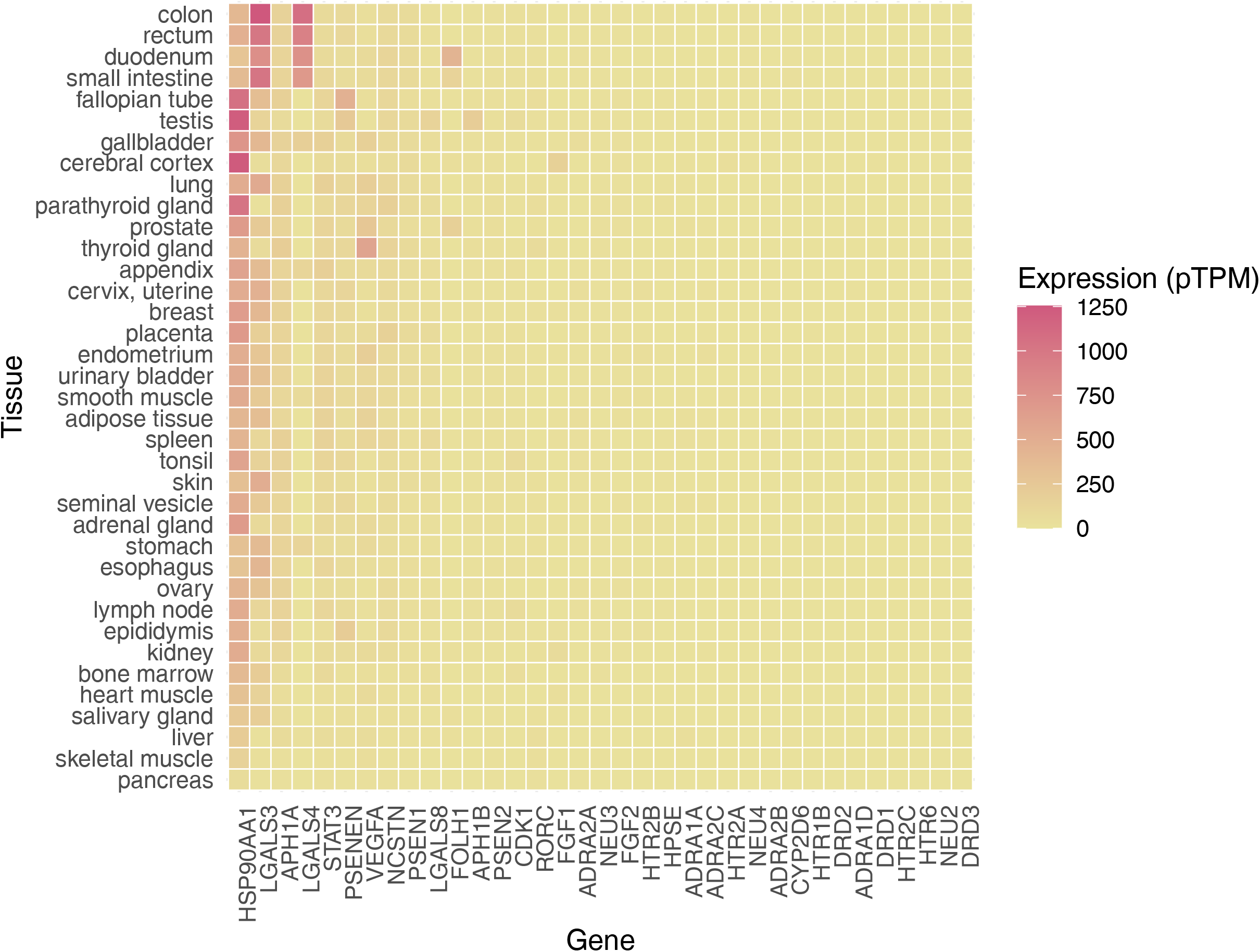
The mRNA transcript expression profile of the predicted trehalose targets in human. The expression profile is categorised based on the pTPM ranging from 0 to 1250.

We also identified subcellular protein expression of the targets of trehalose in human. Since trehalose does not readily permeate the plasma membrane, we focused on proteins which are secreted or expressed on cell junctions, plasma membrane, and secreted vesicles (**figure 4**). We observed that presenilin-1 was expressed on the plasma membrane and is also predicted to be secreted. Gamma-secretase is likely of considerable interest to trehalose safety/bioactivity assessment together with CDK1. Cytochrome P450 2D6 is the only other identified protein expressed on the plasma membrane, however the affinity of trehalose to this target is very low. FGF3 and RORC were located in the vesicles. In addition, we observed a number of proteins expressed in the cell junction, including alpha-1a adrenergic receptor (ADRA1A), alpha-2b adrenergic receptor (ADRA2B), CDK1, dopamine D1 receptor (DRD1), Heat shock protein HSP 90-alpha (HSP90AA1), galectin-3 (LGALS3), nicastrin (NCSTN), and STAT3. All these targets are highly relevant to the assessment of bioactivity of trehalose in pharmaceutical formulations and hence should be integrated into the drug development protocols.

**Figure 4:**
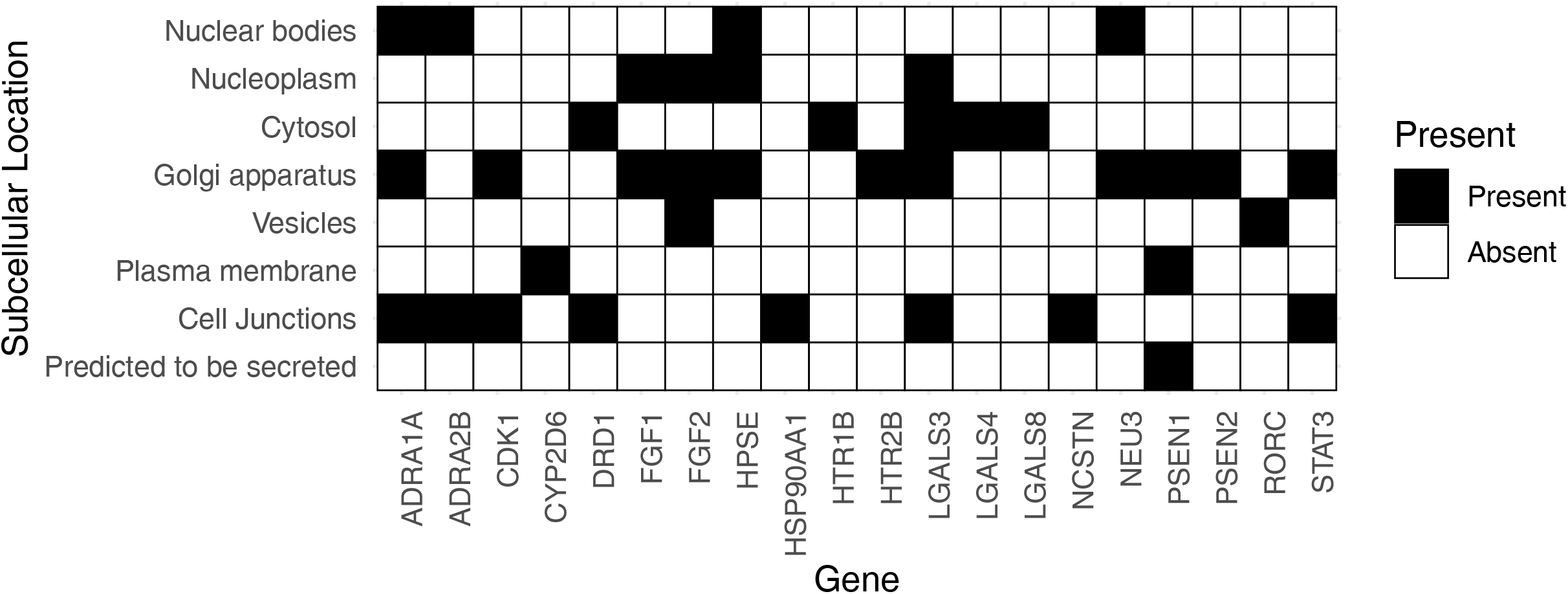
The subcellular localization of the predicted trehalose targets in human.

## Discussion

Trehalose is a Generally Recognized as Safe (GRAS) compound which is widely used as an excipient in food products and more recently in pharmaceutical formulations due to its desirable pharmacokinetic properties. The GRAS label for trehalose is largely based on historical evidence of safety and a limited number of assessments using rodent and rabbit models.^[6–9, 38–40]^ Although excipients are assumed to be pharmacologically inert, this can largely be attributed to a lack of suitable bioassays to quantify their pharmacodynamic effects in parallel to historical evidence of their safety. However, no chemical entity can be considered truly inert, and this especially applies to pharmaceutical excipients which are often used in much higher concentrations than the API. Advancements in bioinformatic and cheminformatic tools address these gaps by making it feasible to objectively assess the pharmacodynamic activity of excipients with selected species specificity.^[41–45]^

This study highlights several pharmacodynamic features of trehalose which are relevant to its use in pharmaceutical formulations. The desirable physicochemical and pharmacokinetic properties of trehalose observed in this study are consistent with several previous *in vivo* studies ^[6, 9, 38, 39, 46, 47]^ which validate its use as a pharmaceutical excipient. Among these desirable excipient properties are its lipophilicity, poor gastrointestinal permeability, slower metabolism, and negligible interference with the liver cytochrome p450 enzymes. Besides these, trehalose is also reported to enhance protein stability, which supports its preferential use in formulating biological therapeutics.^[6–9, 48–50]^ The pharmacodynamic results are consistent with *in vivo* findings, so this study validates the use of *in silico* tools in drug development.

Trehalose is selectively metabolised into monosaccharides by trehalase expressed in the kidney and intestine. The kinetics of this metabolic process are likely slow because the affinity of trehalose to human trehalase was observed to be approximately 66 μM. This observation is consistent with reports of peak trehalase concentration lasting for about 5 hours post intravenous administration in humans which is almost twice the duration typically reported for other disaccharides.^[8, 39, 51]^ We found significant inter-species variability in the affinity of trehalose to trehalase highlight the limitations of rodent studies as models for efficacy analysis of pharmaceutical formulations consisting of trehalose, and perhaps other excipients, too. We observed major expression of trehalase was observed in the kidneys and gastrointestinal system (duodenum and small intestines) which is consistent with previous findings.^[52]^ These observations are critical to screening of all parenteral formulations consisting of trehalose, as such formulations will be preferentially eliminated by renal excretion and therefore such formulations should be critically monitored for its impact on renal physiology.

We observed significant variation in the target classes in human, mouse, and rat. While Family A GPCRs were the most targeted protein class, the remaining target classes considerably varied. The GPCR repertoire varies considerably among mammals, for example the canine GPCR genome is closer to that of humans than it is to mouse and rat.^[53]^ Only 50% of GPCRs have a one-to-one ortholog between human and rat, as well as only 70% between mouse and rat, and the authors suggest that the GPCR repertoire may be more highly diverged than other subsets of the genome.^[54]^ This is reflected in the serotonin receptors, a family of GPCRs, where only one of five receptors were predicted binding targets of trehalose. This further emphasises irrelevance of rodent models in efficacy and safety analysis in preclinical drug discovery and development programs.

Among the many targets of trehalose identified in this study, its affinity for CDK1 was the highest. However, the expression of CDK1 under physiological conditions was observed to be medium to low in human tissues, providing further assurance to the safety of trehalose use in pharmaceutical formulations. Such assurance on safety of trehalose are consistent with several other *in vivo* studies.^[6, 8, 9, 39, 40, 51, 55]^ The number of low affinity targets of trehalose when analysed considering the expression level and subcellular localization of these targets further assure that the impact of trehalose interaction with these targets on systemic physiology will be negligible. However, some of the targets of trehalose identified in this study may become relevant under specific pathological conditions (where the target are upregulated or their sensitivity enhanced), and they should be focused on in the efficacy and safety assessments of trehalose formulations.

In summary, the comparative pharmacodynamic analysis of trehalose highlights the limitations of rodent models, specifically mouse and rat, in translating findings for the development of human therapeutics. The *in silico* analyses performed in this study may provide valuable additions to drug development programs as refinement tools for efficiency and cost optimization. It will be interesting to see if *in silico* comparative pharmacodynamics can be integrated into preclinical programs to make necessary corrections to improve translatability. Without such an approach, rodent model based preclinical programs add limited insight into drug discovery and development at the expense of time and costs.

## Acknowledgement

Research support from University College Dublin-Seed funding/Output Based Research Support Scheme (R19862, 2019) and Stemcology (STGY2917, 2022) is acknowledged.

## References

1. Alhaider, I. A., Mohamed, M. E., Ahmed, K. K. M. and Kumar, A. H. S. Date Palm (Phoenix dactylifera) Fruits as a Potential Cardioprotective Agent: The Role of Circulating Progenitor Cells. Frontiers in Pharmacology, 2017, 8.

2. Saini, A. K., Kumar H S, A. and Sharma, S. S. Preventive and curative effect of edaravone on nerve functions and oxidative stress in experimental diabetic neuropathy. European Journal of Pharmacology, 2007, 568, 164–72.

3. Arun, K. H. S., Kaul, C. L. and Poduri, R. Tempol augments angiotensin II-induced AT2 receptor-mediated relaxation in diabetic rat thoracic aorta. Journal of Hypertension, 2004, 22, 2143–52.

4. Kumar, V., et al. Active pharmaceutical ingredient (API) chemicals: a critical review of current biotechnological approaches. Bioengineered, 2022, 13, 4309–27.

5. Pifferi, G. and Restani, P. The safety of pharmaceutical excipients. Farmaco, 2003, 58, 541–50.

6. Varela Junior, A. S., et al. Trehalose in extenders for cryopreservation of Tambaqui (Colossoma macropomum) sperm. Cryo Letters, 2022, 43, 264–68.

7. Tian, Y., et al. Intradermal Administration of Influenza Vaccine with Trehalose and Pullulan-Based Dissolving Microneedle Arrays. Journal of Pharmaceutical Sciences, 2022, 111, 1070–80.

8. Pupyshev, A. B., Klyushnik, T. P., Akopyan, A. A., Singh, S. K. and Tikhonova, M. A. Disaccharide trehalose in experimental therapies for neurodegenerative disorders: Molecular targets and translational potential. Pharmacological Research, 2022, 183, 106373.

9. Cunha, A., et al. Trehalose-Based Nucleolipids as Nanocarriers for Autophagy Modulation: An In Vitro Study. Pharmaceutics, 2022, 14, 857.

10. Baldrick, P. Pharmaceutical Excipient Development: The Need for Preclinical Guidance. Regulatory Toxicology and Pharmacology, 2000, 32, 210–18.

11. Elbein, A. D., Pan, Y. T., Pastuszak, I. and Carroll, D. New insights on trehalose: A multifunctional molecule. Glycobiology, 2003, 13, 17R–27R.

12. Watanabe, M., Kikawada, T. and Okuda, T. Increase of internal ion concentration triggers trehalose synthesis associated with cryptobiosis in larvae of Polypedilum vanderplanki. Journal of Experimental Biology, 2003, 206, 2281–86.

13. Argüelles, J.-C. Why Can’t Vertebrates Synthesize Trehalose? Journal of molecular evolution, 2014, 79, 111–16.

14. Ishihara, R., et al. Molecular cloning, sequencing and expression of cDNA encoding human trehalase. Gene, 1997, 202, 69–74.

15. Ohtake, S. and Wang, Y. J. Trehalose: Current Use and Future Applications. Journal of Pharmaceutical Sciences, 2011, 100, 2020–53.

16. Guo, N., Puhlev, I., Brown, D. R., Mansbridge, J. and Levine, F. Trehalose expression confers desiccation tolerance on human cells. Nature Biotechnology, 2000, 18, 168–71.

17. Worrall, E. E., Litamoi, J. K., Seck, B. M. and Ayelet, G. Xerovac: An ultra rapid method for the dehydration and preservation of live attenuated Rinderpest and Peste des Petits ruminants vaccines. Vaccine, 2000, 19, 834–39.

18. Bracken, M. B. Why animal studies are often poor predictors of human reactions to exposure. J R Soc Med, 2009, 102, 120–2.

19. Mak, I. W., Evaniew, N. and Ghert, M. Lost in translation: animal models and clinical trials in cancer treatment. Am J Transl Res, 2014, 6, 114–8.

20. Ledford, H. Translational research: 4 ways to fix the clinical trial. Nature, 2011, 477, 526–8.

21. Daina, A., Michielin, O. and Zoete, V. SwissTargetPrediction: updated data and new features for efficient prediction of protein targets of small molecules. Nucleic acids research, 2019, 47, W357–W64.

22. Manchukonda, B. and Kumar, A. H. Network profiling of hepatocellular carcinoma targets for evidence based pharmacological approach to improve clinical efficacy. Biology, Engineering, Medicine and Science Reports, 2022, 8(1), 11–15.

23. Kumar, A. H. S. Pharmacological targets of Asundexian relevant to its therapeutic efficacy in treating cardiovascular diseases. Research Square, 2022.

24. Khosravi, Z., Kaliaperumal, C. and Kumar, A. Analysing the role of SERPINE1 network in the pathogenesis of human glioblastoma. bioRxiv, 2022.

25. Sagar, V. and Kumar, A. H. Efficacy of natural compounds from Tinospora cordifolia against SARS-CoV-2 protease, surface glycoprotein and RNA polymerase. Biology, Engineering, Medicine and Science Reports, 2020, 6, 06–08.

26. Kumar, A. PTPRC, KDM5C, GABBR1 and HDAC1 are the major targets of valproic acid in regulation of its anticonvulsant pharmacological effects. Biology, Engineering, Medicine and Science Reports, 2022, 8(2), 28–32.

27. Khosravi Z and AH., K. Pharmacognosy and pharmacology of Calotropis gigantea for discovery of anticancer therapeutics. Pharmacognosy Magazine., 2021, Apr 1;17(6):, 123–27.

28. Kumar, A. H. and Sharma, V. Acetamido-Propanoic Acid Derived Compounds as Protease Inhibitors to Target Coronaviruses. Biology, Engineering, Medicine and Science Reports, 2019, 5, 20–22.

29. Hadley, W. Ggplot2: Elegrant graphics for data analysis. (Springer, 2016).

30. Kumar, A. H. Molecular docking of natural compounds from tulsi (Ocimum sanctum) and neem (Azadirachta indica) against SARS-CoV-2 protein targets. Biology, Engineering, Medicine and Science Reports, 2020, 6, 11–13.

31. Goothy, S. S. K. and Kumar, A. H. Network proteins of angiotensin-converting enzyme 2 but not angiotensin-converting enzyme 2 itself are host cell receptors for SARS-Coronavirus-2 attachment. Biology, Engineering, Medicine and Science Reports, 2020, 6, 01–05.

32. Santamaría, D., et al. Cdk1 is sufficient to drive the mammalian cell cycle. Nature, 2007, 448, 811–15.

33. Brunkan, A. L. and Goate, A. M. Presenilin function and gamma-secretase activity. Journal of Neurochemistry, 2005, 93, 769–92.

34. Barondes, S. H., Cooper, D. N. W., Gitt, M. A. and Leffler, H. Galectins. Structure and function of a large family of animal lectins. Journal of Biological Chemistry, 1994, 269, 20807–10.

35. Seto, J. T. and Rott, R. Functional significance of sialidase during influenza virus multiplication. Virology, 1966, 30, 731–37.

36. Mackenzie, F. and Ruhrberg, C. Diverse roles for VEGF-A in the nervous system. Development (Cambridge, England), 2012, 139, 1371–80.

37. Imada, K. and Leonard, W. J. The Jak-STAT pathway. Molecular Immunology, 2000, 37, 1–11.

38. Pan, S., et al. Trehalose ameliorates autophagy dysregulation in aged cortex and acts as an exercise mimetic to delay brain aging in elderly mice. Food Science and Human Wellness, 2022, 11, 1036–44.

39. Richards, A. B., et al. Trehalose: A review of properties, history of use and human tolerance, and results of multiple safety studies. Food and Chemical Toxicology, 2002, 40, 871–98.

40. Liu, M., et al. Multiple toxicity studies of trehalose in mice by intragastric administration. Food Chemistry, 2013, 136, 485–90.

41. Kumar, D. A. H. S. and Sharma, V. Acetamido-Propanoic Acid Derived Compounds as Protease Inhibitors to Target Coronaviruses. Biology, Engineering, Medicine and Science Reports, 2019, 5, 20–22.

42. Goothy, S. S. K. and Kumar, A. H. S. Network Proteins of Angiotensin-converting Enzyme 2 but Not Angiotensin-converting Enzyme 2 itself are Host Cell Receptors for SARS-Coronavirus-2 Attachment. Biology, Engineering, Medicine and Science Reports, 2020, 6, 01–05.

43. Kumar, A. H. S. Pharmacological Targets of Asundexian Relevant to its Therapeutic Efficacy in Treating Cardiovascular Diseases. Biology, Engineering, Medicine and Science Reports, 2022, 8, 24–27.

44. Kumar, A. H. S. Network Pharmacology Analysis of Orally Bioavailable SARS-CoV-2 Protease Inhibitor Shows Synergistic Targets to Improve Clinical Efficacy. Biology, Engineering, Medicine and Science Reports, 2021, 7, 21–24.

45. Kumar, A. H. S. Pharmacology of Chloroquine: Potential Mechanism of Action against Coronavirus. Biology, Engineering, Medicine and Science Reports, 2020, 6, 09–10.

46. Tanaka, M., et al. Trehalose alleviates polyglutamine-mediated pathology in a mouse model of Huntington disease. Nature Medicine, 2004, 10, 148–54.

47. Higashiyama, T. Novel functions and applications of trehalose. Pure and Applied Chemistry, 2002, 74, 1263–69.

48. Olsson, C., Jansson, H. and Swenson, J. The Role of Trehalose for the Stabilization of Proteins. The Journal of Physical Chemistry B, 2016, 120, 4723–31.

49. Kaushik, J. K. and Bhat, R. Why Is Trehalose an Exceptional Protein Stabilizer? An Analysis of the Thermal Stability of Proteins in the Presence of the Compatible Osmolyte Trehalose. Journal of Biological Chemistry, 2003, 278, 26458–65.

50. Jovanović, N., et al. Distinct effects of sucrose and trehalose on protein stability during supercritical fluid drying and freeze-drying. European Journal of Pharmaceutical Sciences, 2006, 27, 336–45.

51. Argov, Z., Vornovitsky, H., Blumen, S. and Caraco, Y. First Human Use of High Dose IV Trehalose: Safety, Tolerability and Pharmacokinetic Results from the Oculopharyngeal Muscular Dystrophy (OPMD) Therapy Trial (P7.068). Neurology, 2015, 84.

52. Oesterreicher, T. J., Markesich, D. C. and Henning, S. J. Cloning, characterization and mapping of the mouse trehalase (Treh) gene. Gene, 2001, 270, 211–20.

53. Haitina, T., Fredriksson, R., Foord, S. M., Schiöth, H. B. and Gloriam, D. E. The G protein-coupled receptor subset of the dog genome is more similar to that in humans than rodents. BMC Genomics, 2009, 10, 24.

54. Gloriam, D. E., Fredriksson, R. and Schiöth, H. B. The G protein-coupled receptor subset of the rat genome. BMC Genomics, 2007, 8, 338.

55. Tang, K.-K., Liu, X.-Y., Wang, Z.-Y., Qu, K.-C. and Fan, R.-F. Trehalose alleviates cadmium-induced brain damage by ameliorating oxidative stress, autophagy inhibition, and apoptosis. Metallomics: Integrated Biometal Science, 2019, 11, 2043–51.

